# Nitrogen conservation, conserved: 46 million years of N-recycling by the core symbionts of turtle ants

**DOI:** 10.1101/185314

**Authors:** Yi Hu, Jon G. Sanders, Piotr Łukasik, Catherine L. D’Amelio, John S. Millar, David R. Vann, Yemin Lan, Justin A. Newton, Mark Schotanus, John T. Wertz, Daniel J. C. Kronauer, Naomi E. Pierce, Corrie S. Moreau, Philipp Engel, Jacob A. Russell

## Abstract

Nitrogen acquisition is a major challenge for herbivorous animals, and the repeated origins of herbivory across the ants have raised expectations that nutritional symbionts have shaped their diversification. Direct evidence for N-provisioning by internally housed symbionts is rare in animals; among the ants, it has been documented for just one lineage. In this study we dissect functional contributions by bacteria from a conserved, multi-partite gut symbiosis in herbivorous *Cephalotes* ants through *in vivo* experiments, (meta)genomics, and *in vitro* assays. Gut bacteria recycle urea, and likely uric acid, using recycled N to synthesize essential amino acids that are acquired by hosts in substantial quantities. Specialized core symbionts of 17 studied *Cephalotes* species encode the pathways directing these activities, and several recycle N *in vitro*. These findings point to a highly efficient N-economy, and a nutritional mutualism preserved for millions of years through the derived behaviors and gut anatomy of *Cephalotes* ants.

**Category:** Biological Sciences-Evolution

## Introduction

Nitrogen (N) is a key component of living cells and a major constituent of the nucleic acids and proteins directing their structure and function. Like primary producers^1^, herbivorous animals face the challenge of obtaining sufficient N in a world with limited accessible N, suffering specifically due to the low N content of their preferred foods^2^. The prevalence of herbivory is, hence, a testament to the many adaptations for sufficient N-acquisition. Occasionally featured within these adaptive repertoires are internally housed, symbiotic microbes. Insects provide several examples of such symbioses, with disparate herbivore taxa co-opting symbiont N-metabolism for their own benefit^3, 4, 5, 6^. Such tactics are not employed by all insect herbivores^7^, however, and few studies have quantified symbiont contributions to host N-budgets^8, 9, 10, 11, 12^.

Ants comprise a diverse insect group with a broad suite of diets. Typically viewed as predators or omnivores, several ants are functional herbivores, with isotopic N-ratios overlapping those of known herbivorous insects^13, 14^. While occasionally obtaining N from tended sap-feeding insects, most are considered plant canopy foragers, scavenging for foods such as extrafloral nectar, pollen, fungi, vertebrate waste, and plant wound secretions^14^. Quantities of usable and essential N in such foods are limiting^15, 16^. Hence, the repeated origins of functional herbivory provide a useful natural experiment, enabling tests for symbiotic correlates of N-limited diets. The concentration of specialized bacteria within herbivorous ant taxa suggests such a correlation^17, 18^. But N-provisioning by internally housed symbionts has only been documented for carpenter ants, whose intracellular *Blochmannia* provide them with amino acids made from recycled N^19^.

Herbivorous cephalotines (i.e. *Cephalotes* and *Procryptocerus*) and ants from other herbivore genera (i.e. *Tetraponera* and *Dolichoderus*) exhibit hallmarks of a symbiotic syndrome distinct from that in the Camponotini. Large, modified guts with prodigious quantities of extracellular gut bacteria make up one defining feature^20, 21, 22, 23, 24^. Also characteristic are the oral-anal trophallaxis events transmitting symbionts between siblings^21, 25, 26, 27^ and the domination of gut communities by host-specific bacteria^17, 28, 29^. Such symbiotic “hotspots” stand out in relation to several ant taxa, which show comparatively low investment in symbiosis^18, 23^.

N-provisioning by bacterial symbionts in these ants has been hypothesized as a mechanism for their success in a seemingly marginal dietary niche^14, 17^. To investigate this, we focus on the turtle ants of the genus *Cephalotes* (**Fig. 1**). With ~115 described species^30^, stable isotopes place these arboreal ants low on the food chain^14, 31^. Workers are canopy foragers, consuming extrafloral nectar and insect honeydew, fungi, pollen and leaf exudates^32, 33, 34^. *Cephalotes* also consume mammalian urine and bird feces, excreta with large quantities of waste N accessible only through the aid of microbes. Given this, the remarkably conserved gut microbiomes of cephalotines^28, 35^ are proposed as an adaptation for their N-poor and N-inaccessible diets. Here we measure symbiont N-provisioning in *Cephalotes varians* and gene content within the gut microbiomes of 17 *Cephalotes* species (**Table S1**), describing symbiont N-metabolism across 46 million years of evolutionary history.

**Figure 1:**
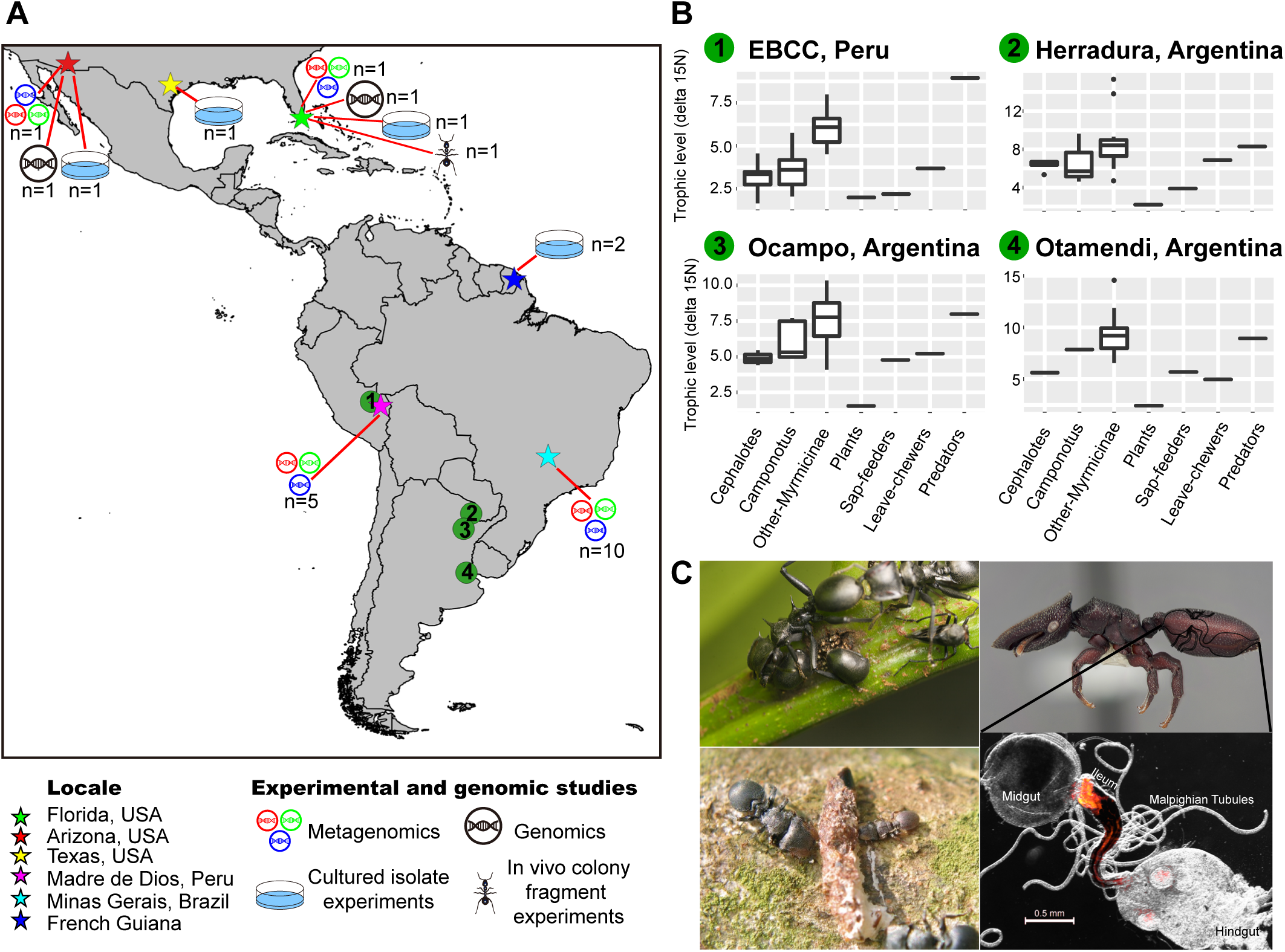
Ecology of *Cephalotes* ants and origins of specimens used in our study. A) Map showing sampling locales for cephalotines (*Cephalotes* and *Procryptocerus*) used in this study (stars), the activities they were used for, along with sample size (i.e. # of species). Numbered circles show sites of ant sampling in two prior studies^14,31^, from which stable nitrogen isotope data were extracted and plotted here. B) Nitrogen isotope ratios obtained from *Cephalotes* ants, other ants in the Myrmicinae, and *Camponotus* ants, hosts of known N-recycling bacteria. High ratios of ^15^N/^14^N in organismal tissues suggest a higher placement on the food chain. Panels show results from different locales, with each illustrating N isotope ratios for plants, known herbivores, and known predators. C) *C. atratus* workers tend honeydew-producing, ant-mimicking membracids (upper left). *C. eduarduli* and *C. maculatus* (smaller worker) feeding on bird droppings (lower left). Soldier caste of *C. varians* with an outlined digestive tract (upper right). A FISH microscopy image of a digestive tract from a *Cephalotes* worker is shown at lower right. Note the large bacterial mass in the ileum near the midgut-ileum junction, the site where N-wastes are emptied via Malpighian tubules.

## Results

### Gut bacteria of turtle ants do not fix N_2_

Atmospheric N_2_-fixation is executed by bacterial symbionts of some invertebrates^36, 37, 38, 39, 40^, and prior detection of nitrogenase genes in ants^17, 29^ has led to the proposal that symbiotic bacteria fix nitrogen for their hosts. To test this, three *Cephalotes varians* colonies were subjected to acetylene reduction assays within hours of field capture. In three separate experiments no ethylene was produced within test tubes containing acetylene-exposed ants (**Table S2**), arguing against active N-fixation.

### Symbiont-upgrading of dietary amino acids has minimal impact on workers’ N-budgets

Based on precedents from intracellular symbionts of insects^12, 19, 41^, we then tested whether gut bacteria could upgrade dietary nitrogen compounds, transforming non-essential or inaccessible N compounds into essential amino acids that are acquired by hosts. Our efforts focused on glutamate, an important pre-cursor in the synthesis of many amino acids. *Cephalotes varians* workers from three colonies were reared on artificial diets^42^ varying in the presence/absence of antibiotics and the presence/absence of heavy isotope labeled glutamate. ^13^C or ^15^N were used to label glutamate across our two separate experiments. Heavy isotope enrichment in the free amino acid pools from worker hemolymph, assessed via GC-MS (**Table S3**), allowed us to quantify symbiont glutamate upgrading and provisioning back to hosts.

Antibiotic treatment successfully suppressed gut microbial load in this and all below experiments (**Fig. S1**), and workers survived treatments at rates sufficient for subsequent data generation (**Fig. S2**). In addition, ants clearly absorbed nutrients from the administered diets, as hemolymph glutamate pools showed 4-7% enrichment for heavy isotopes on the heavy vs. light isotope diets in the absence of antibiotics (p=0.0033 ^15^N vs. ^14^N diet; p=0.0018 ^13^C vs. ^12^C diet). Yet, ant acquisition of symbiont-processed C and N from dietary glutamate was minimal. For instance, on the ^13^C-glutamate diet, antibiotic treatment reduced the fraction of heavy isotope-bearing isoleucine (p=0.0147), leucine (p=0.0004), threonine (p=0.0029), and tyrosine (p=0.0169) in worker hemolymph (**Fig. S3**), relative to estimates on this same diet without antibiotics. But effect sizes for each amino acid were small, with changes of just 1.3-2.6% in ants with suppressed microbiota. On the ^15^N-glutamate diet (**Fig. S4**), only phenylalanine (p=0.045) showed heavy isotope enrichment in untreated vs. antibiotic treated workers, again, with small effect size (2.9%).

### Symbionts recycle, then upgrade dietary N-waste; hosts acquire this N

*Cephalotes* ants consume bird droppings and are attracted to mammalian urine. In addition, most of their symbionts colonize the ileum^25^, the gut compartment where Malpighian tubules deliver nitrogenous wastes (**Fig. 1**). Insects lack the capacity to convert their dominant nitrogenous wastes—uric acid or urea—into usable forms of N. It has, thus, been posited that ant-associated gut symbionts recycle N waste, further converting this N into essential amino acids that are acquired by hosts^21^. To test this, we implemented an experiment similar to our dietary N-upgrading assessments (above), using ^15^N isotope labeled urea instead of glutamate. After consuming diets with heavy urea, 15 of the 16 detectable amino acids in *C. varians* hemolymph were enriched for the heavy isotope signal when compared to diets with light (^14^N) urea (i.e. all but asparagine; significant p-value range: 1.92E-14 - 0.0210) (**Fig. 2**; **Table S3**). On the ^15^N diet, antibiotic treatment strongly reduced the heavy isotope signal in these same 15 amino acids (significant p-values ranged from 1.92E-14 – 0.0410), directly implicating bacteria in the use of diet-derived N-waste. The impact of bacterial metabolism on worker N-budgets was substantial, with 15-36% enrichment of heavy essential amino acids in hemolymph of symbiotic, versus aposymbiotic, ants within five weeks on the experimental diet.

**Figure 2:**
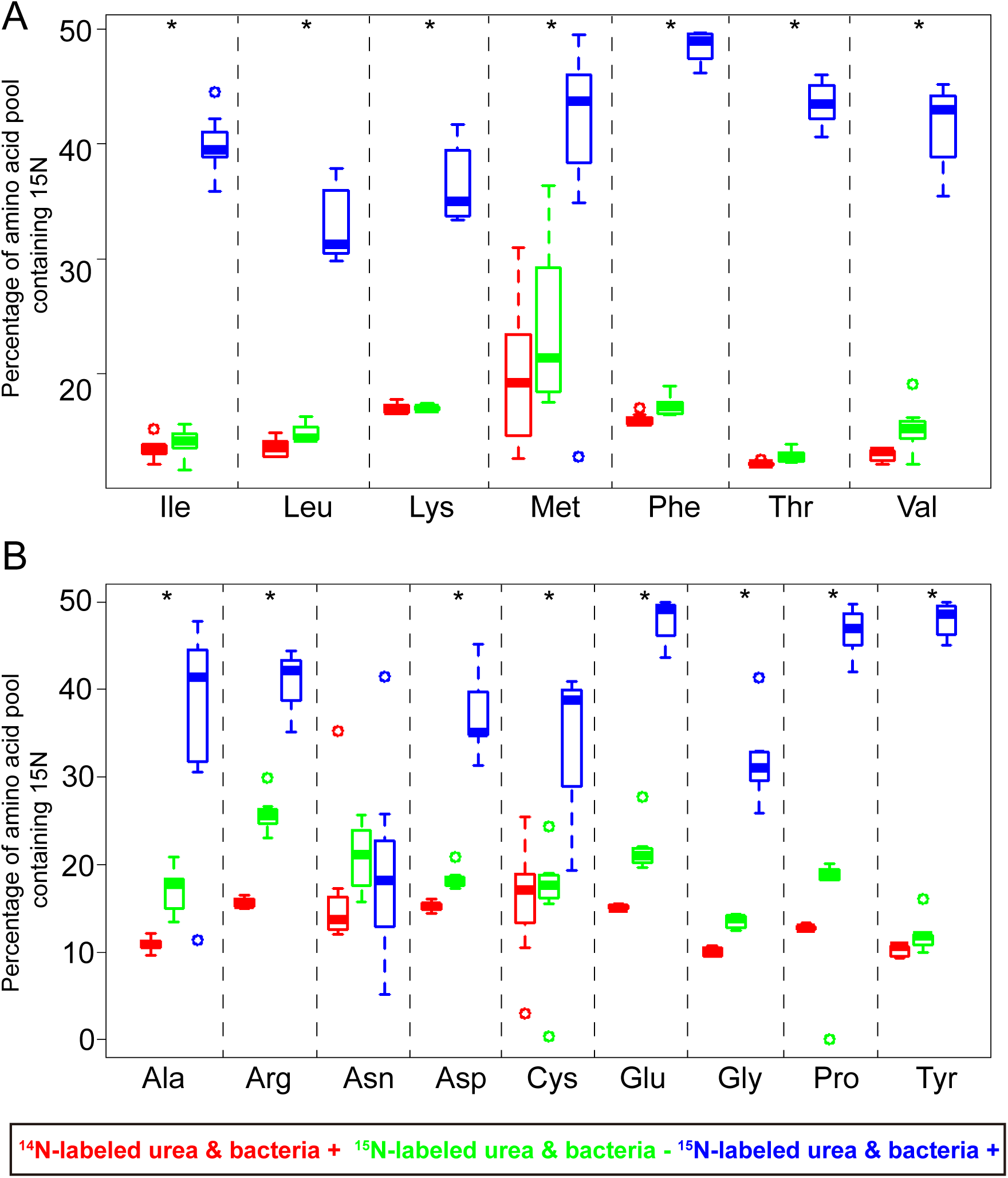
Symbiont removal reduces proportions of free, ^15^N-labeled amino acids in hemolymph of *Cephalotes varians* workers consuming ^15^N-labeled urea. (A) Essential, and (B) non-essential amino acids in ant hemolymph measured through GC-MS. Asterisks indicate that ^15^N in essential amino acids from ants consuming ^15^N-labeled urea (blue) was significantly higher than that in antibiotic-treated ants on this same diet (green) and in those consuming diets with unlabeled urea (red).

### Metagenomic analyses: strong taxonomic conservation

To address the mechanisms behind symbiont N-recycling and upgrading, we used shotgun Illumina HiSeq sequencing to characterize microbiomes. Eighteen sequence libraries were generated across seventeen *Cephalotes* species collected from four locales (**Fig. 1**; **Table S1**). Two of these came from our experimental model *C. varians*. Shotgun sequencing efforts yielded median values of 32,706,498 reads and 143.625 Mbp of assembled scaffolds per library (**Table S4**). The median N50 for scaffold length was 1106.5 bp.

A prior study suggested that >95% of the *Cephalotes* gut community is comprised of core symbionts from host-specific clades^43^. To assess whether the dominant bacteria sampled here came from such specialized groups we extracted 16S rRNA fragments ≥200bp from each metagenome library. Top BLASTn hits were downloaded for each sequence, and jointly used in a maximum likelihood phylogenetic analysis. In the resulting tree (**Fig. S5**), 94.4% of our 335 *Cephalotes* symbiont sequences grouped within 10 cephalotine clades that included sequences from prior *in vivo* studies. Inferences on metagenome content have, hence, been made using partial genomes from the dominant, specialized core taxa.

Classification of assembled scaffolds took place using USEARCH comparisons against public reference genomes in IMG and the KEGG database (see Materials & Methods, and Supplementary Methods for more detail). Results from these analyses paralleled our 16S rRNA-based discoveries of a highly conserved core microbiome (**Fig. 3**). In all metagenomes, *Cephaloticoccus* symbionts^44^ from the Opitutales were the most dominant, with scaffolds from these bacteria typically forming a single “cloud” differentiated from others by depth of coverage and %GC content. Xanthomonadales scaffolds were ubiquitous and typically abundant, with multiple scaffold clouds often evidencing co-existence of distinct strains, with up to ~10% average %GC divergence. Less abundant, though still ubiquitous across metagenomes were clouds of scaffolds from the Pseudomonadales, Burkholderiales, and Rhizobiales. Multiple scaffold clouds at differing depths of coverage were consistent with multiple strain co-existence, for each taxon, within most microbiomes. Unlike these core groups, Flavobacteriales, Sphingobacteriales, and Campylobacterales were common but not ubiquitous. For instance, presence/absence calls for N-metabolism genes (**Table S5**) suggested the absence or extreme rarity of these symbionts in several metagenomes.

**Figure 3:**
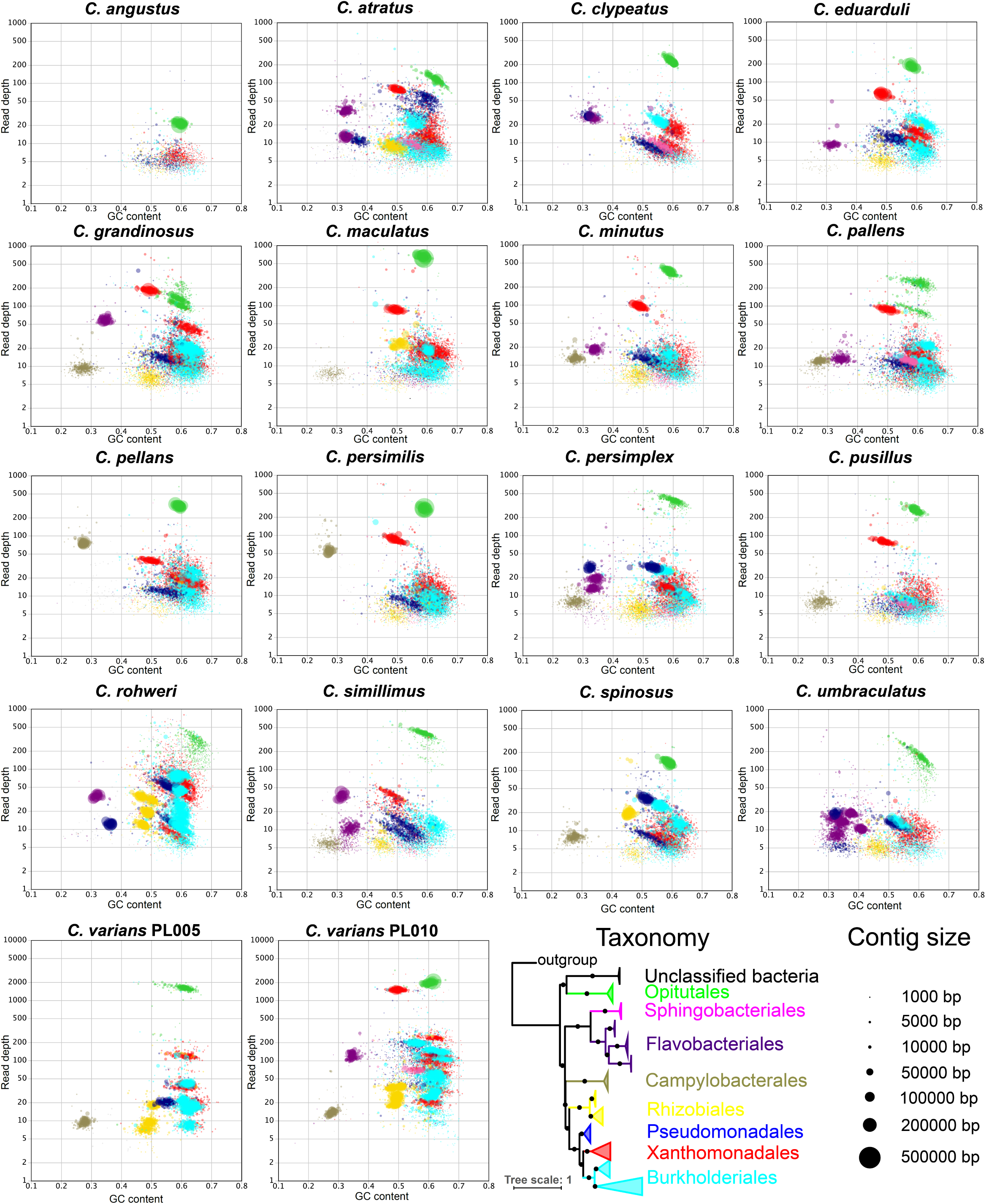
Taxon-annotated GC-coverage plots for 18 *Cephalotes* metagenomes, reveals taxonomic conservation. Assembled scaffolds in each metagenome are plotted based on their %GC content (x-axis) and their depth of sequencing coverage (y-axis, log scale). Bacterial genomes vary in %GC genome content and core symbionts show variable abundance; these plots, thus, illustrate the existence of numerous dominant symbiont strains in *Cephalotes* worker guts. Phylogeny at lower right, based on 16S rRNA sequences from our metagenomes, identify the *Cephalotes*-specific clades from which nearly all of our sequence data have been obtained. Colors on the phylogeny match those in the blob plots, illustrating the taxa to which scaffolds were assigned. Circle size reveals scaffold length. Not shown here are scaffolds binning to Hymenoptera or to unclassified organisms.

### Metagenomic analyses: urease is ubiquitous, N_2_ fixation is absent

Consistent with our acetylene reduction experiments using *C. varians*, IMG/ER based annotation recovered no N-fixation genes in any of the 18 metagenome libraries. This absence encompassed genes encoding the molybdenum-containing nitrogenase system (i.e. *nifD, nifH, nifK*), and those from the iron-only (*anfD*, *anfG*, *anfH*, *anf*K) and vanadium-containing (*vnfD*, *vnfG*, *vnfH*, *vnfK*) systems (**Table S5**). Together, our experiments and metagenomics suggest that prior observations of *nifH* genes in *Cephalotes* workers arose from detection of rare or contaminant bacteria^17^ or from a portion of the gut not included in the present study (exclusively the midgut, ileum, and rectum).

Matching our discovery of symbiotic N-recycling in *C. varians* were findings of *ureA*, *ureB*, and *ureC* genes in both *C. varians* metagenomes and in those of the 16 additional *Cephalotes* species (**Figs. 4**, **S6**, **S7**; **Table S5**). The presence of complete gene sets for the core protein subunits of the urease enzyme in all sampled microbiomes suggests that symbionts from most *Cephalotes* species can make ammonia from N-waste urea. Taxonomic classification for urease gene-encoding scaffolds suggested that abundant *Cephaloticoccus* symbionts (order: Opitutales) encoded all three core urease genes. Complete copies of each gene were found on a single Opitutales-assigned scaffold within 15 of 18 metagenomes. Genes encoding the urease accessory proteins (*ureF, ureG*, and *ureH*) were often found on these same scaffolds, with a strong trend of conserved architecture for this gene cluster (**Fig. S6**).

**Figure 4:**
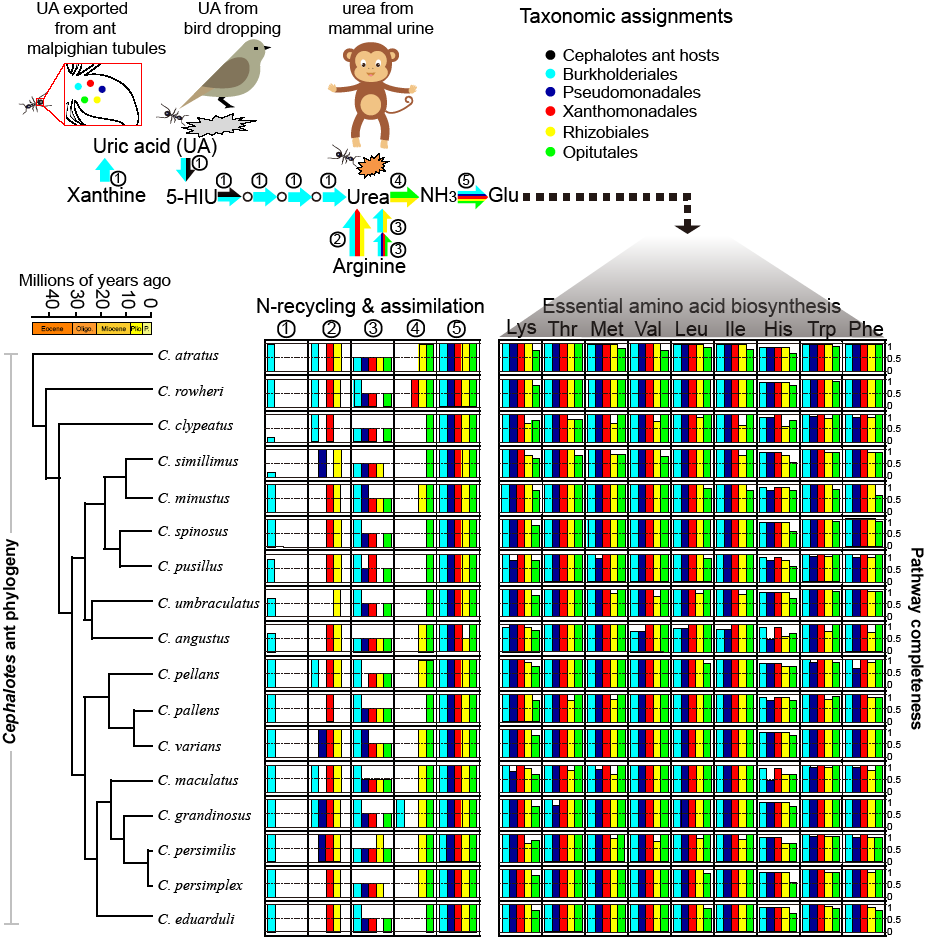
Pathways for N-waste recycling and amino acid biosynthesis and their distributions across core gut symbionts from 17 *Cephalotes* species. Various symbiotic gut bacteria convert N-wastes into ammonia, incorporate ammonia into glutamate, and synthesize essential amino acids. Upper panel shows sources of the N-wastes uric acid (bacterial metabolism via xanthine degradation, bird droppings, host ant waste-metabolism via Malpighian tubule delivery) and urea (mammalian urine, uric acid metabolism, and arginine metabolism). Arrows in this panel are colored to reflect taxonomy of the core *Cephalotes*-specific microbes participating in these steps in multiple metagenomes. Numbers near arrows link particular pathways to bar graphs (below), which in turn plot pathway completeness (i.e. proportion of all genes present) for the dominant core taxa in each metagenome. At left on the lower panel below is the phylogeny of *Cephalotes* species used for metagenomics including a chronogram dating divergence events in these species’ history^30^.

Urease genes were occasionally assigned to other bacteria (**Fig. 4**; **Table S5**), suggesting that more than one symbiont can participate in this recycling function. Notable were cases from *C. rohweri* (Xanthomonadales), *C. grandinosus* (Burkholderiales), and *C. eduarduli* (unclassified Bacteria), which hosted additional bacteria encoding complete sets of urease core and accessory proteins. In these cases, urease genes mapped to single scaffolds with identical gene order to that seen for *Cephaloticoccus* (**Fig. S6**). Urease function was also inferred for Rhizobiales bacteria in several *Cephalotes* species. Rhizobiales-assigned scaffolds encoding urease genes differed from those of *Cephaloticoccus* with respect to gene order, the presence of the *ureJ* accessory gene, and the existence of a gene fusion between *ureA* and *ureB* (**Fig. S6**).

A maximum likelihood phylogenetic analysis of UreC proteins encoded by the sampled microbiomes identified two distinct *Cephalotes*-specific lineages (**Fig. S8**). The first (bootstrap support = 99%) consisted of Rhizobiales-assigned UreC proteins, with relatedness to homologs from various families in the Rhizobiales. The second (bootstrap support = 93%) was comprised of proteins assigned to Opitutales, Burkholderiales, Xanthomonadales, and unclassified Bacteria, showing relatedness to homologs from bacteria in the Rhodocyclales (Betaproteobacteria).

### Metagenomic analyses: ammonia assimilation and amino acid synthesis genes found across numerous core taxa

The above results provide genetic mechanisms to explain symbiont-mediated urea recycling in *C. varians*, suggesting a broad distribution for this function across the *Cepalotes* genus. Also necessary to explain our experiments are: 1. symbiont genes to assimilate the ammonia made from urea degradation, and 2. symbiont genes using this assimilated N to synthesize amino acids. Assessment of our metagenomes met these expectations in *C. varians* and all 16 other host species. But in contrast to our findings for a small number of urea recyclers, genes involved in these processes assigned to all core symbiont taxa, suggesting extensiove metabolic overlap.

Within all metagenomes, numerous taxa encoded complete gene sets for ammonia assimilation (e.g. **Figs. 4**, **S7, S9, S10; Table S5**). Similarly, gene sets for the synthesis of each essential and non-essential amino acid were complete in all metagenomes. With the exception of histidine synthesis, complete only for Campylobacterales, each amino acid biosynthesis pathway was complete within multiple bacterial taxa (**Figs. 4**, **S6**, **S7, S9, S10; Table S5**). Xanthomonadales and Burkholderiales bins (outside of *C. angustus*) encoded all genes to synthesize the remaining eight essential amino acids. This paralleled findings for Opitutales, Rhizobiales, and Pseudomonadales: the former typically showed an incomplete pathway for lysine, while the latter two often seemingly lacked a single gene for methionine biosynthesis. Sphingobacteriales and Flavobacteriales lacked required genes in the lysine and methionine pathways. And pathways for threonine, valine, leucine, isoleucine, tryptophan, and phenylalanine were occasionally missing all or most genes within the Flavobacteriales. Many of the core taxa also possessed complete gene sets for the synthesis of non-essential amino acids.

### Metagenomic analyses: uric acid synthesis and degradation, and other means of urea production

Uric acid is a major waste product of many insects. This compound is also found in bird excreta, a common *Cephalotes* food. To analyze capacities to recycle N from uric acid we examined gene content in the pathway converting this compound into urea. Scaffolds assigning to Hymenoptera, thus likely originating from *Cephalotes* genomes, often contained a subset of genes involved in uric acid degradation, including one encoding the canonical uricase enzyme *(uaZ)* and one encoding breakdown of 5-HIU *(uraH;* **Table S5**; **Fig. S11**).

Beyond those scaffolds, Burkholderiales bacteria were implicated in uricase function in all but two metagenomes (**Figs. 3**, **4**, **S6, S7; Table S5**). First, the *puuD* uricase homolog was detected in 14 metagenomes. Encoding a membrane-associated form of this enzyme, with a C-terminal cytochrome c domain^45^, this gene was found on Burkholderiales-assigned scaffolds in all cases where detected. Genes encoding the remaining steps in the uric acid degradation pathway (i.e. 5-HIU→OHCU→allantoin→allantoate→urea) also classified to Burkholderiales (**Fig. S6**). In total, our analyses suggested complete gene sets for this pathway and taxon in 13 out of 18 metagenomes (**Table S5**; **Figs. 4**, **S7**). Maximum likelihood phylogenies of the bacterially encoded PuuD and UraH proteins revealed monophyly of homologs from *Cephalotes*-associated Burkholderiales (bootstrap support = 98% for PuuD and 76% for UraH; **Fig. S12**). Lineages in both trees showed relatedness to homologous proteins from free-living Burkholderiales and other Proteobacteria.

Genes synthesizing urea from uric acid mapped to numerous scaffolds across several metagenomes. However, in seven libraries, they mapped to just one or two Burkholderiales-assigned scaffolds. Synteny was conserved in these cases, and such scaffolds also possessed additional genes encoding the subunits of xanthine dehydrogenase (**Fig. S6**), an enzyme converting xanthine into uric acid. Core symbionts appear to produce xanthine via purine recycling, as genes for guanine deaminase enzymes (**Fig. S13**) classified to Burkholderiales in 16 metagenomes (**Table S5**). Adenine deaminase enzymes were similarly encoded by bacteria, namely Rhizobiales, across 10 metagenomes.

Further analyses of our metagenomes revealed that bacteria aside from Burkholderiales can produce urea through other mechanisms (**Fig. 4**; **Table S5**). For instance, across most hosts, microbes from the Burkholderiales, Rhizobiales, Xanthomonadales, and/or Pseudomonadales possessed arginase genes, catalyzing a reaction that converts arginine to urea and ornithine. In several metagenomes, arginase genes also binned to Hymenoptera, suggesting their presence in *Cephalotes* genomes. Genes for a separate, two-step pathway converting arginine to urea (**Fig. S14**) were present in most metagenomes. But only Burkholderiales encoded both steps.

### *Refining symbiont role assignments: fine scale metagenome binning, cultured core symbiont genomes, and* in vitro *N-recycling assays*

Multiple strains for many of the aforementioned core taxa often co-exist within a single gut community^43^. So despite pathway “completeness” assessed at the level of host order, it remains unclear whether individual symbiont strains encode complete pathways for key aspects of N-metabolism. We addressed this through genome sequencing of cultured symbiont strains from five of the eight core bacterial taxa across two host species (i.e. *C. varians* and *C. rohweri*). The 14 strains prioritized for sequencing were chosen based on 16S rRNA gene identity (or near identity) in comparison to core symbionts previously sampled through culture-independent techniques (**Fig. S5**). Alignments of *C. varians* isolates to scaffolds from conspecific metagenomes (**Fig. S15**) indicates that these strains or very close relatives are present *in vivo*, supporting the relevance of *in vitro* findings from these strains to the natural gut community..

Highlights of this work (**Fig. 5**; **Supplementary Results; Table S7**) included the discovery of a Burkholderiales strain (Cv33a) with a capacity to convert uric acid into urea. This strain lacked urease genes, but three cultured symbionts encoded all genes necessary for urease function, including *Cephaloticoccus* isolates from *C. varians* (Cv41) and *C. rohweri* (Cag34) and a Xanthomonadales symbiont from *C. rohweri* (Cag60). Thirteen out of fourteen isolates encoded the glutamate dehydrogenase gene (*gdhA*) converting ammonia into glutamate, and most encoded complete pathways for synthesizing most amino acids. As expected from metagenomic analyses, all genomes lacked nitrogenase genes.

**Figure 5:**
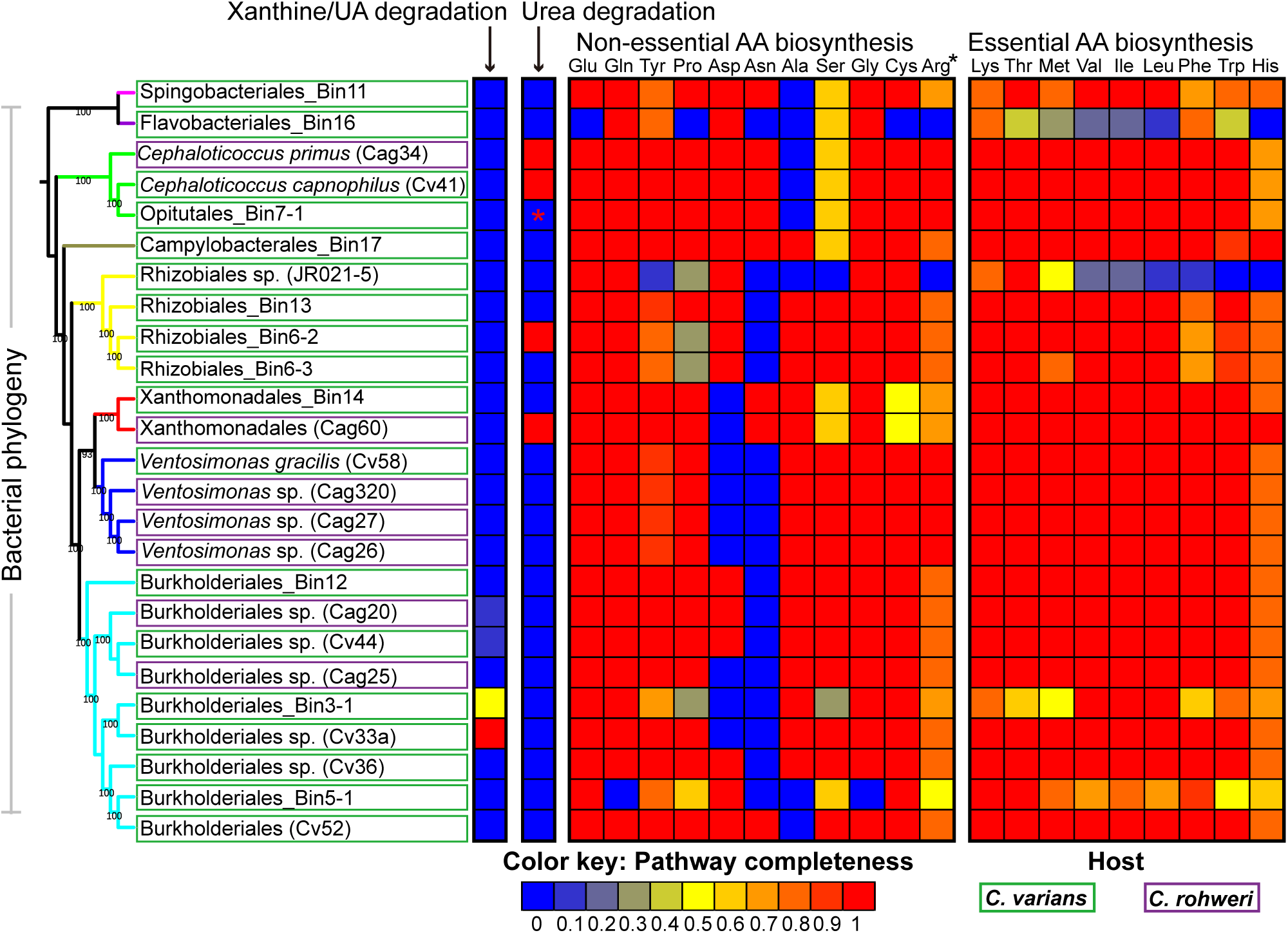
Core symbiont strains possess complete or near complete pathways for N-recycling and amino acid biosynthesis. Heatmap illustrates the proportion of genes present from each N-metabolic pathway across distinct symbiont strains. Coding capacities for strains were inferred from 14 fully sequenced cultured isolate genomes (symbionts from *C. varians* and *C. rohweri*) and 11 draft genomes (assembled from *C. varians* colony PL010 metagenome; identified by the term “Bin” within their names). The maximum likelihood phylogeny of symbiotic bacteria on the left was inferred using an alignment of amino acids encoded by seven phylogenetic marker genes obtained from symbiont genomes, and branch colors are used to illustrate distinct bacterial orders. Red asterisk for urea recycling in the *Cephaloticoccus*-like Opitutales bin (7-1) indicates that urease genes from the PL010 metagenome binned to Opitutales, but not to the draft genome for the dominant strain. When combined with the likely presence of just one Opitutales strain within the PL010 microbiome, it is likely that a completely assembled genome would encode all urease genes. The black asterisk next to Arg denotes that inferences on pathway completeness for arginine biosynthesis were based on the pathway starting with glutamate (see Fig. S9), as opposed to other metabolic mechanisms.

The fastidious nature of some symbionts limited our ability to infer strain functions for common core taxa. Insights for these groups were gained through draft genome assembly from our best sampled metagenome (i.e. *C. varians* colony PL010) using the Anvi’o platform (version 1.2.3)^46^ in conjunction with the CONCOCT differential coverage-based binning program^47^. The 11 near complete draft genomes, where ≥87% of universal single copy genes were detected, spanned seven of our eight core symbiont taxa (**Fig. 5**; **Tables S8**, **S9**, **S10**). Gene content analyses supported findings from metagenomics and cultured isolate genomes (**Supplementary Results**). In short, the dominant symbiont strains individually encoded up to 17 complete amino acid biosynthesis pathways. Incomplete pathways were often missing just one to two genes. Nearly all draft strain genomes showed capacities to assimilate ammonia into glutamate. N-recycling pathways appeared incomplete within predicted N-recycling Burkholderiales and *Cephaloticoccus* strains. This suggests the occasional absence of this function from these taxa (but see **Fig. 6**) or, possibly, incomplete genome assembly due to challenges of scaffold binning (**Supplementary Results**).

**Figure 6:**
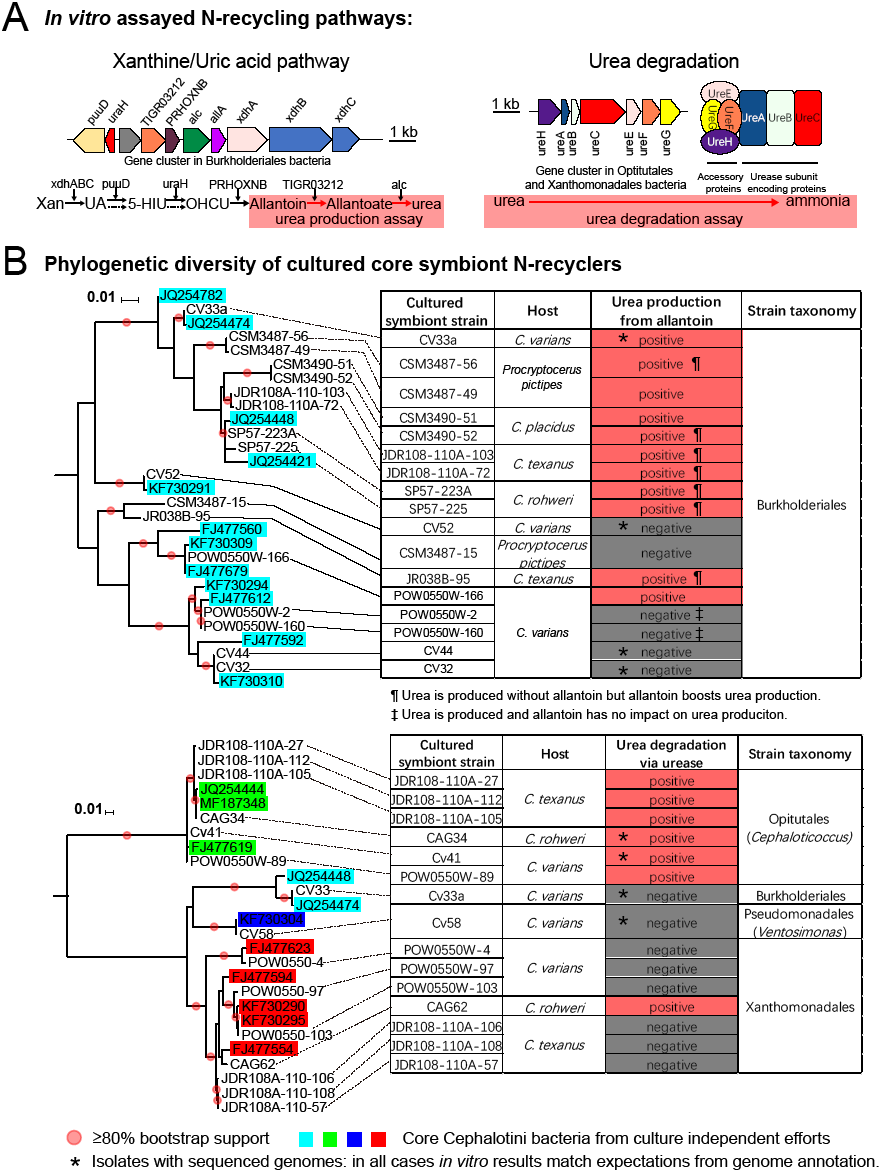
A limited range of cultured core symbiont strains recycle N in vitro. Results summarize findings from symbionts of five cephalotine ants, including four species from the genus *Cephalotes* and one from its sister genus P*rocryptocerus.* A) Genes and pathways used by specialized gut symbionts to recycle the N-wastes uric acid and urea. The architecture for clusters of symbiont genes encoding the involved enzymes are illustrated. Red boxes within these pathways represent the metabolic steps assayed for panel B. B) Shown at left are phylogenies of cultured symbionts subject to metabolic assays in vitro and their closest relatives in the NCBI database, which are enclosed within taxon-specific colored boxes. Most cultured isolates had 16S rRNA sequences that were identical or most closely related to one obtained from a *Cephalotes* ant through culture-independent means. Such isolates also showed high identity to abundantly represented scaffolds from our metagenomes (**Fig. S15**). Nodes for cultured symbionts are connected to relevant rows within data tables, where the results of assays for urea production (from allantoin—part of the uric acid pathway) and urea degradation (to ammonia) assays are illustrated. Asterisks highlight isolates with a sequenced genome; for each of these, in vitro results matched expectations derived from gene content. Additional symbols used in the urea production table indicate whether allantoin boosted urea production and whether production was completely allantoin-dependent. Urea production without complete allanotin-dependence is likely to stem from arginine metabolism.

To test whether genetic signatures reflect actual N-recycling capacities, and to study the conservation of this role within key taxa, we performed a series of *in vitro* assays, expanding the number of targeted cephalotine species. Urease activity was qualitatively assessed by the generation of ammonia in the presence of urea, and we obtained positive results for seven of fifteen tested isolates (**Fig. 6**). All *Cephaloticoccus* (Opitutales) were positive, as was one of six Xanthomonadales isolates. Results for four isolates with sequenced genomes accurately reflected predictions from the presence/absence of urease genes.

Production of urea from allantoin served as a proxy for activity of the xanthine/uric acid pathway (**Fig. 6**). Urea was produced from allantoin for 11 of the 17 assayed Burkholderiales isolates (**Table S11**), suggesting function for at least part of this pathway. Coding capacity from the five isolates with sequenced genomes accurately predicted the results of these assays.

In summary, genomic inferences on N-recycling seem to accurately reflect the metabolism of core symbionts. And importantly, the phylogenetic placement of strains with *in vitro* assay data reveal sporadic distributions of N-recycling, with notable enrichment in two clades (*Cephaloticoccus* and a specific, unnamed lineage of Burkholderiales) suggesting long-standing roles in the efficient use of N by the *Cephalotes* holobiont.

## Discussion

Our findings show that ancient, specialized gut bacteria of *Cephalotes* ants recycle waste N acquired through the diet (urea) and, likely, through ant waste metabolism (urea and uric acid). Workers acquire large amounts of symbiont-recycled N in the form of essential and non-essential amino acids. Symbionts encode genes to derive their own uric acid and urea, suggesting a third potential origin for the influx of waste N into this system. Across a broad range of *Cephalotes* species, gene content for N-metabolism varies little within core taxa and N-recycling roles appear conserved within specific symbiont lineages. These findings depict an efficient N-economy retained across 46 million years of *Cephalotes* evolution. They also support the hypothesis that this multi-partite gut microbiome plays an adaptive role within an N-poor dietary niche.

The magnitude of symbiont contributions to host N-budgets has rarely been calculated. But, findings from wood-feeding termites implicate N-fixing bacteria in providing up to 60% of the N in termite colonies^10^. Measures from the leaf-cutter ant system suggest that N-fixing bacterial symbionts provide 45-61% of the fungus garden’s N-supply^48^. Our estimate in *C. varians* that 15-36% of the free amino acid pool was derived from symbiont-recycled N, within five weeks of feeding, was notable, though not directly comparable to either of these estimates. Reduced survival of antibiotic-treated workers, on diets where urea was the only source of N (**Fig. S2**), do however suggest the importance of symbiont N-metabolism in adults. A similar importance of bacteria was previously suggested for *Cephalotes atratus*^49^. In carpenter ants *Blochmannia* have noticeable impacts on worker performance, larval and pupal development, and colony growth; and the detriments of *Blochmannia* removal can be partially alleviated by the addition of essential amino acids to the diets of aposymbiotic ants^19^. While the impacts of *Cephalotes* worker microbiomes on larvae have not been measured, adult N-stores are implicated in larval nourishment for several ants^50^. These results suggest a large potential for symbionts of adults to shape performance at all stages within the colony.

The ubiquity of N-recycling *Blochmannia* across the Camponotini^19, 51^ combine with our findings to support the hypothesized importance of nutritional symbionts in canopy-dwelling, herbivorous ants^14^. A trend of “convergent associations”^52^ has, thus, emerged: canopy foraging for N-poor or N-inaccessible foods has evolved separately in association with unrelated, yet functionally similar symbionts. Future work on other ant herbivores and their conserved symbionts^23, 29^ will assess the generality of such functional convergence. Also of interest will be studies of symbiont-independent strategies for navigating N-poor diets^7, 53^.

The conserved nature of symbiont community composition across cephalotines is remarkable compared to patterns for many arthropods^54, 55, 56^, adding to a trend across eusocial insects. Within the termites, for instance, many core symbionts hail from host-specific lineages, revealing ancient, specialized relationships^3^. Among the corbiculate bees, some relationships with gut symbionts date back to 80 million years^57^. But even for these hosts, occasional symbiont turnover takes place—in association with dietary shifts, for termites^58^, and among major phylogenetic divisions in bees^57^.

Evolved behaviors have likely preserved partner fidelity in these groups. Among eusocial bees, symbiont transfer takes place within the hive, through a combination of trophallaxis, coprophagy, or contact with nest materials^59, 60, 61^. Termite siblings transmit symbionts through oral-anal trophallaxis^62, 63^. A similar mode of passage has been noted for *Cephalotes* and *Procryptocerus* ants and for other ant herbivores as well^21, 25, 26, 27^. Of likely further importance for cephalotines is a fine-mesh filter, enveloping the proventriculus, which can bar the passage of particles as small as 0.2 μM beyond the crop. This filter develops shortly after young adults solicit trophallactic symbiont transfers^25^. Symbionts acquired during early adulthood will, thus, be sealed off within the midgut, ileum, and rectum, with minimal opportunities for subsequent colonization by additional, ingested microbes. These dual drivers of partner fidelity^64^ may collectively explain the preservation of an ancient nutritional mutualism and sustained exploitation of N-poor foods by successful canopy herbivores.

## Materials & Methods

### Collections and experimental assays

Details on ant collections and the uses of ants from particular locales are presented in **Fig. 1** and **Table S1.** For many of these protocols, additional details can be found in the **Supplementary Methods**.

Experiments on live ants were performed on *Cephalotes varians* colony fragments collected from the Florida Keys. Acetylene reduction assays were used to assess the capacity for N-fixation. To achieve this, we incubated adult workers (and also, in some instances, queens, larvae, and pupae) in air-tight syringe chambers loaded with acetylene very shortly after collection in the field. After incubation, samples were analyzed with a gas chromatography-flame ionization detector to quantify levels of acetylene and ethylene.

Controlled lab experiments were performed to quantify microbial contributions toward N-upgrading of non-essential dietary amino acids and, separately, N-recycling, and subsequent upgrading, of dietary N-waste. Adult workers from each experimental colony (n=3 colonies per each of three experiments) were divided into three groups with equal number. In the first treatment, workers were fed antibiotics to suppress or eliminate their gut bacteria for three weeks. After this time, workers transitioned to the trial period where they were continuously fed antibiotics in addition to a diet of glutamate (with ^15^N or ^13^C) or a diet with urea containing ^15^N. Feeding for this trial period lasted four to five additional weeks. For the second and third treatment groups, bacterial communities were not disrupted. Diets for workers in these groups were identical to those of treatment group one, save for antibiotics, during the three week pre-trial period. For the four to five week trial period, workers from the second treatment group were fed on the aforementioned heavy-isotope diets; those from the third group were fed otherwise identical diets in which glutamate or urea consisted of standard isotope ratios (i.e. biased toward lighter isotopes).

During the trial period we recorded worker mortality, noting an elevation in the ^15^N urea feeding group treated with antibiotics, but not in antibiotic-treated workers from the one examined glutamate experiment (**Fig. S2**). Efficacies of antibiotic treatments were quantified via qPCR on bacterial 16S rRNA genes and, for a subset of specimens, amplicon sequencing of 16S rRNA (**Fig. S1; Table S14**). Worker hemolymph was harvested at the end of the four to five week trial, from three to ten surviving workers per replicate. Hemolymph was then pooled, used for amino acid derivitization, and subjected to GC-MS to quantify proportions of free amino acids containing the heavy isotopes (see details in **Table S12**).

### Metagenomics

Adult workers were dissected under a light microscope using fine forceps, with removal of gut tissues from each dissected *Cephalotes* worker. DNA extractions were performed on ten single guts for ten workers from each of two colonies for *C. varians* or on pools of guts from ten workers in a single colony for each of the remaining *Cephalotes* species. Separate extractions from *C. varians* siblings were then pooled within colonies. The resulting two DNA samples and the 16 samples from other *Cephalotes* species were then used for metagenomic library preparation. After size selection, adapter ligation, amplification, and clustering, samples were sequenced (2x100bp or 2x150 paired end reads) on an Illumina HiSeq2500 machine. Sequences were trimmed for quality, with removal of adapter sequences after de-multiplexing. Assembly of reads from individual libraries then proceeded using a variety of k-values with the IDBA-UD metagenomic assembler. Scaffolds were uploaded to the Integrated Microbial Genomes with Microbiome Samples Expert Review (IMG/M-ER) website^65^. Classification in IMG/MER proceeded based on USEARCH similarity against all public reference genomes in IMG and the KEGG database. Due to some incorrect scaffold assignments (to errant bacterial taxa not known from *Cephalotes*, such as Rhodocyclales), seven genomes from cultured bacterial isolates belonging to core *Cephalotes*-associated taxa were used to obtain more accurate information of phylogenetic binning. IMG/M-ER was used to annotate gene content from our scaffolds and taxonomic bins. Based on these annotations, we focused on N-metabolism, using KEGG^66^ and Metacyc^67^ as guides to manually construct degradation pathways for N-waste products and biosynthetic pathways for amino acids. We examined the completeness of the N-waste degradation pathways based on Fig. 6A and the completeness of the amino acid biosynthetic pathways based on Figs. S7 and S9 across 18 metagenomes, 14 isolate genomes and 11 draft genomes (as described below).

Homologs from N-recycling pathways and 16S rRNA genes were extracted from each dataset and used in phylogenetic analyses with closely related homologs from the NCBI database. To further aid in understanding taxonomic composition and to illustrate depth of coverage for the taxa in our libraries, we generated “blob plots” based on read mapping to classified scaffolds using BWA^68^ and modified scripts from a prior publication^69^. These graphs showed the depth of coverage for each scaffold in relation to our classifications, along with the %GC content, a taxonomically conserved genomic signature that further aided us in our efforts to visualize the diversity of symbionts within microbiomes (**Fig. 3**)

### Fine-scale metagenomic binning to generate draft symbiont genomes

To improve the assignment of metabolic capabilities to individual symbiont strains we used the Anvi’o metagenome visualization and annotation pipeline (version 1.2.3)^46^ in conjunction with the CONCOCT differential coverage-based binning program^47^. In doing so, we binned assembled scaffolds from the metagenomic datasets of *C. varians* colony PL010—the library with best symbiont coverage—into draft genomes of symbiont strains. Reconstruction of N-metabolic pathways was then performed to comprehend the range of metabolic capabilities of individual symbionts.

### *Genomics and* in vitro *assays on cultured bacterial isolates*

Gut tissues from *Cephalotes* and *Procryptocerus* worker ants were dissected and macerated. Contents were then plated on tryptic soy agar plates, and plates were incubated at 25°C under an atmosphere of normal air supplemented with 1% carbon dioxide in a CO_2_-controlled water-jacketed incubator. After colony sub-cloning, pure isolate cultures were maintained under these same conditions on the aforementioned plates or in tryptic soy broth. DNA extracted from these cultures was subsequently used for 16S rRNA PCRs to compare isolates to bacteria previously sampled through culture-independent studies. Isolates from *C. varians* and *C. rohweri* (both previously well-studied through culture-independent means) were prioritized for genome sequencing when their 16S rRNA sequences were identical or nearly identical to those of from prior *in vivo* studies. Extracted bacterial DNA was used for library preparation and Illumina or PacBio SMRT sequencing. Assembled genomes were uploaded to IMG/ER for annotation, with N-metabolism pathway reconstruction and extraction of genes for phylogenetics occurring as described above. Alignments of isolate genomes to metagenomes were done with Icarus^70^ as implemented in MetaQuast^71^ and visualized in Circos^72^

A subset of cultured isolates was subjected to assays to detect ammonia production from urea. Several were also tested to determine whether allantoin, a derivative of uric acid breakdown, could be used to synthesize urea. Methodological details on these assays are described in the **Supplementary Methods**. As described above for genome sequencing prioritization, strains prioritized for assays were those deemed highly related to specialized core *Cephalotes* symbionts.

### Fluorescence In Situ Hybridization

The fixed, dissected gut of an adult worker from *Cephalotes* sp. JGS2370 was hybridized with a set of eubacterial probes labeled with AlexaFluor488 and AlexaFluor555 and imaged following the protocol in **SI Appendix, Supplementary Methods** on a Leica M165FC microscope. Fluorescent microphotographs taken using blue and green excitation filters were then merged with a photograph taken under white light.

## Acknowledgments

We thank Ryuichi Koga for assistance with Fluorescence In Situ Hybridization and Amit Basu for help with amino acid derivitization. YH’s dissertation committee members Sue Kilham, Shivanthi Anandan, Mike O’Connor and Sean O’Donnell provided useful suggestions on statistics and experimental design. Dr Pamela Plantinga provided advice on statistical analyses for *in vitro* symbiont assays. This study was funded by NSF grant #s 1050360 and 1442144 to JAR, NSF grant #1442316 to CSM, and NSF grant #1442156 to JTW. Funding was also provided by SNFS grant IZK0Z3_164213 to YH and PE, and by SNFS grant 31003A_160345 to PE.

